# Inference of distribution of fitness effects and proportion of adaptive substitutions from polymorphism data

**DOI:** 10.1101/062216

**Authors:** Paula Tataru, Maéva Mollion, Sylvain Glemin, Thomas Bataillon

**Author notes:** Bioinformatics Research Centre, Aarhus University, C.F. Møllers Alle 8, Aarhus C 8000, Denmark.

## Abstract

The distribution of fitness effects (DFE) encompasses deleterious, neutral and beneficial mutations. It conditions the evolutionary trajectory of populations, as well as the rate of adaptive molecular evolution (*α*). Inference of DFE and *α* from patterns of polymorphism (SFS) and divergence data has been a longstanding goal of evolutionary genetics. A widespread assumption shared by numerous methods developed so far to infer DFE and *α* from such data is that beneficial mutations contribute only negligibly to the polymorphism data. Hence, a DFE comprising only deleterious mutations tends to be estimated from SFS data, and *α* is only predicted by contrasting the SFS with divergence data from an outgroup. Here, we develop a hierarchical probabilistic framework that extends on previous methods and also can infer DFE and *α* from polymorphism data alone. We use extensive simulations to examine the performance of our method. We show that both a full DFE, comprising both deleterious and beneficial mutations, and *α* can be inferred without resorting to divergence data. We demonstrate that inference of DFE from polymorphism data alone can in fact provide more reliable estimates, as it does not rely on strong assumptions about a shared DFE between the outgroup and ingroup species used to obtain the SFS and divergence data. We also show that not accounting for the contribution of beneficial mutations to polymorphism data leads to substantially biased estimates of the DFE and *α*. We illustrate these points using our newly developed framework, while also comparing to one of the most widely used inference methods available.

---

New mutations are the ultimate source of heritable variation. The fitness properties of new mutations determine the possible evolutionary trajectories a population can follow (Bataillon and Bailey 2014). For instance, supply rate and fitness effects of beneficial mutations determine the expected rate of adaptation of a population (Lourenço *et al.* 2011), while deleterious mutations condition the expected drift load of a population (Kimura et al.1963). Even a few beneficial mutations with large effects can quickly move a population towards its fitness optimum, while the fitness can be reduced through the accumulation of multiple deleterious mutations with small effects that occasionally escape selection. Genome-wide rates and effects of new mutations influence, among others, the evolutionary advantage of sex (Otto and Lenormand 2002), the expected degree of parallel evolution (Chevin et al. 2010b), the maintenance of variation on quantitative traits (Hill 2010), and the evolutionary potential and capacity of populations to respond to novel environments (Chevin *et al.* 2010a; Hoffmann and Sgrò 2011).

Effects of new mutations on fitness are typically modeled as independent draws from an underlying distribution of fitness effects (hereafter DFE) which, in principle, spans deleterious, neutral and beneficial mutations. Lately, there has also been considerable focus on estimation of the DFE of new non-synonymous mutations, and learn more about factors governing the rate of adaptive molecular evolution, commonly defined as the proportion of fixed adaptive mutations among all non-synonymous substitutions, and often denoted *α*. Therefore, inferring the DFE, both from experimental (Bataillon and Bailey 2014; Bataillon *et al.* 2011; Jacquier *et al.* 2013; Halligan and Keightley 2009; Sousa *et al.* 2011), but also from polymorphism and divergence data (Eyre-Walker *et al.* 2006; Keightley and Eyre-Walker 2007; Boyko *et al.* 2008; Eyre-Walker and Keightley 2009; Keightley and Eyre-Walker 2012; Galtier 2016), has been a longstanding goal of evolutionary genetics.

The McDonald-Kreitman test (McDonald *et al.* 1991) was one of the first attempts to use DNA data to measure the amount of selection experienced by genes. It compares the amount of variation (counts of nucleotide polymorphism) within a species (ingroup) to the variation between species (measured by divergence counts between sequences from the ingroup and an outgroup). The test parses and contrasts the amount of variation found at the synonymous and non-synonymous sites, where the synonymous sites are assumed to be neutrally evolving sites. Smith and Eyre-Walker (2002) further developed this test to also infer the amount of purifying selection, defined as the proportion of strongly deleterious mutations, and *α* (see also Welch (2006) for a maximum likelihood approach). Building on the Poisson Random Field (PRF) theory (Sawyer and Hartl 1992; Sethupathy and Hannenhalli 2008) and arising as extensions to the classical McDonald-Kreitman test, a series of methods have been developed to not only characterize the amount of selection, but also the DFE (Bustamante *et al.* 2003; Piganeau and Eyre-Walker 2003; Eyre-Walker *et al.* 2006; Keightley and Eyre-Walker 2007; Boyko *et al.* 2008; Keightley and Eyre-Walker 2010; Gronau *et al.* 2013; Kousathanas and Keightley 2013; Racimo and Schraiber 2014), and then used it as a building block to estimate *α* (Loewe *et al.* 2006; Eyre-Walker and Keightley 2009; Schneider *et al.* 2011; Keightley and Eyre-Walker 2012; Galtier 2016).

Assuming that sites are independent, that new mutations follow a Poisson process and always occur at new sites, these methods then model the observed variation using a Poisson distribution. The variation within the ingroup is given through counts of the site frequency spectrum (SFS), whose mean in each entry of the SFS is calculated as a function of the DFE and other parameters. Selection is assumed to be weak (s ≪ 1, but note that 4*N*_*e*_*s* can still be large) and the DFE to be constant in time and the same in both the ingroup and outgroup.

Additionally, in order to disentangle selection from demography and other forces (Nielsen 2005), and in the spirit of the McDonald-Kreitman test, the sequenced sites are divided into two classes of neutrally evolving and selected sites. The DFE is then inferred by contrasting the SFS counts for the neutral and selected sites, by assuming that such forces equally affect the two classes.

Ideally, a full demographic model should be jointly inferred with the DFE parameters from the data. However, this can be computationally very demanding and instead a simplified demography is often assumed, where a single population size change is allowed (Keightley and Eyre-Walker 2007; Eyre-Walker and Keightley 2009; Kousathanas and Keightley 2013), or a somewhat more complex model is inferred (Boyko *et al.* 2008). Alternatively, the explicit inference of demography can be avoided altogether by introducing a series of nuisance parameters thataccount for the demography and sampling effects. These parameters account for distortions of the polymorphism counts relative to neutral expectations in an equilibrium Wright-Fisher population (Eyre-Walker *et al.* 2006; Galtier 2016). An added benefit is that controlling for demography effects (either explicitly or through nuisance parameters) can also remove bias caused by linkage (Kousathanas and Keightley 2013; Messer and Petrov 2012). The approach of Eyre-Walker *et al.* (2006) can potentially be more robust for estimating a DFE than putting a lot of faith in a simplified demographic scenario.

The proportion of adaptive substitutions, *α*, is typically obtained as a ratio between an estimate of the number of adaptive substitutions and the observed selected divergence counts (Eyre-Walker and Keightley 2009; Loewe *et al.* 2006; Keightley and Eyre-Walker 2012; Galtier 2016). The number of adaptive substitutions is calculated by subtracting, from the observed divergence counts at selected sites, the expected counts accrued by fixation of deleterious and neutral mutations. These expected counts are calculated from an inferred DFE of deleterious mutations (henceforth denoted deleterious DFE). The deleterious DFE is inferred from the SFS data under the assumption that all SNPs at selected sites are only deleterious. Therefore this approach for estimating *α* heavily relies on the assumption that the ingroup and outgroup species share the same DFE-or more accurately, the same distribution of scaled selection coefficients *S* =4*N*_*e*_*s*. Unfortunately, this assumption of invariance might not often be met in practice, because the DFE might change, or simply because it is unlikely that both ingroup and outgroup evolved with the same population size.

There has been great focus on developing methods inferring a deleterious DFE from polymorphism data alone (Keightley and Eyre-Walker 2007; Kousathanas and Keightley 2013; Eyre-Walker *et al.* 2006; Racimo and Schraiber 2014). These methods rely on a crucial assumption: beneficial mutations contribute negligibly to polymorphism (SFS counts) and therefore are not modeled for this type of data. The reasoning behind this is that strongly selected beneficial mutations will fixate very quickly and that “at most an advantageous mutation will contribute twice as much heterozygosity during its lifetime as a neutral variant” (Smith and Eyre-Walker 2002). This assumption is backed by one study (Keightley and Eyre-Walker 2010) discussed in more details in the *Results and Discussion* section. While some DFE methods do model a full DFE (encompassing both deleterious and beneficial mutations) (Bustamante *et al.* 2003; Piganeau and Eyre-Walker 2003; Boyko *et al.* 2008; Schneider *et al.* 2011; Gronau *et al.* 2013; Galtier 2016), the majority of them do not estimate *α*.

Here, we develop a hierarchical probabilistic model that combines and extends previous methods, and that can infer both the full DFE and *α* from polymorphism data alone. We use our method and perform extensive simulations to investigate different aspects of the inference quality. We show that the assumption that beneficial mutations make negligible contribution to SFS data is unfounded and that a full DFE can also be inferred reliably from polymorphism data alone. Using the estimated full DFE, we show how *α* can be inferred without relying on divergence data. Performing inference on polymorphism data alone proves more adequate when assumptions regarding the outgroup evolution (for example, that the scaled DFE is shared between the ingroup and outgroup) are not likely to be met. We also demonstrate that when the contribution of beneficial mutations to SFS data is ignored, both the inferred deleterious DFE and *α* can be heavily biased. We compare our method and illustrate the resulting bias using the most widely used inference method, dfe-alpha (Keightley and Eyre-Walker 2007; Eyre-Walker and Keightley 2009; Schneider *et al.* 2011; Keightley and Eyre-Walker 2012). We also investigate the impact on inference of misidentification of ancestral state.

## Hierarchical model for inference of DFE and *α*

In this study, we build on several of the methods using PRF theory to build a hierarchical model to infer, via maximum likelihood, the DFE from polymorphism (site frequency spectrum, SFS) and divergence counts. Our hierarchical model is combining and extending different features from different approaches. Figure 1 shows a schematic of the data and the model. We offer below a summary of the assumptions and theory underlying our approach. Further details on the likelihood function, its implementation and numerical optimization can be found in the Supplemental Material.

**Figure 1.**
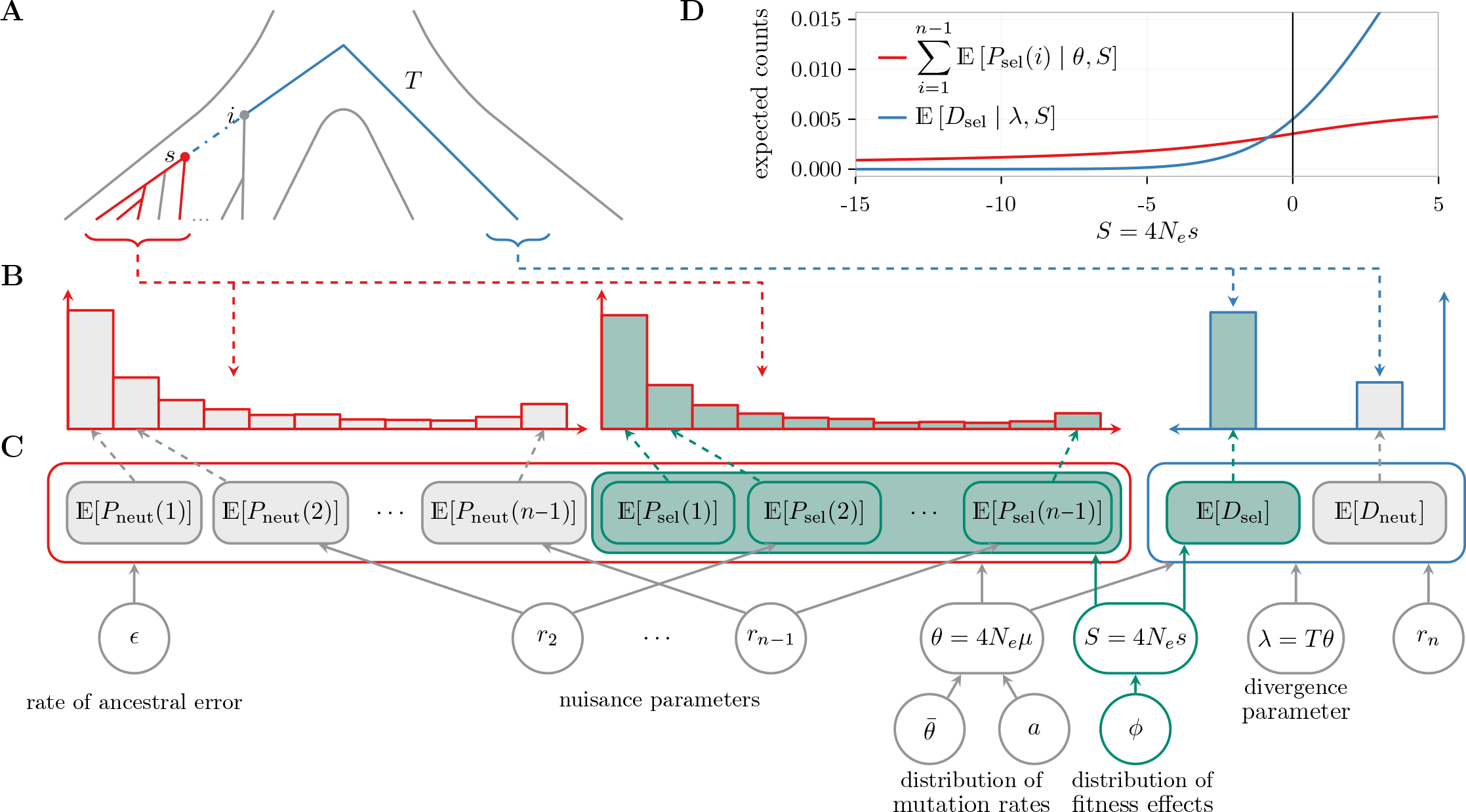
Schematic of data and model. Throughout the figure, gray and green filling indicates sites that are assumed to be evolving neutrally or potentially under selection, respectively, while red and blue outline indicates polymorphism and divergence data (expectations), respectively. (A) The history and coalescent tree of two populations: the ingroup (on the left side), for which polymorphism data is collected, and the outgroup (on the right side), for which divergence counts are obtained. A total of *n* sequences are sampled from the ingroup (marked in red), with the most recent common ancestor (MRCA) found at *s*. The MRCA of the whole ingroup population is found at *i*. From the outgroup we typically have access to one sequence (marked in blue). The total evolutionary time between *s* and the sampled outgroup sequence can be divided into the time from *s* to *i* (blue dot-dash line) and *T*, the time from *i* to the sampled outgroup sequence (blue full line). (B) Site frequency spectrum and divergence counts (*p*_*z*_(*i*) and *d*_*z*_, with **z** ∈ {neut, sel} and 1 ≤ *i* < *n*). (C) Expected counts (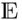[*P*_*z*_(*i*)] and 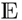[*D*_*z*_], with *z* ∈ {neut, sel} and 1 ≤ *i* | *n*), model parameters and relations between parameters, expectations and data. (D) Expectations as a function of *S*, for θ = 0.001 and λ = 0.005.

### Notations and assumptions

The data is divided into sites that are assumed to be either sites that evolve neutrally (henceforth marked by the subscript neut), or sites that bear mutations with fitness consequences and for which the DFE is estimated (henceforth marked by the subscript sel). Let the observed SFS be given through *p*_*z*_(*i*), where *p*_*z*_(*i*) is the count of polymorphic sites that contain the derived allele *i* times, 1 ≤ *i* < *n*, and *l*_*z*_ the total number of sites surveyed, where *n* is the sample size and *z* ∈ {neut, sel}. We denote by *P*_*z*_ (*i*) the corresponding random variable per site, defined as the random number of sites that contain the derived allele *i* times, normalized by *l*_*z*_. From the PRF theory, *p*_*z*_(*i*) follows a Poisson distribution with mean *l*_z_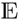 [*P*_*z*_(*i*) | *θ*, *ϕ*], where *θ* = 4*N*_*e*_*μ* is the scaled mutation rate per site per generation, and *ϕ* is a parametric DFE (Figure 1B and C) that will be specified later in the *Results and Discussion* section. Here, we assume additive selection and we define the selection coefficient *s* as the difference in fitness between the heterozygote for the derived allele and the homozygote for the ancestral allele, leading to fitness of 1, 1 + *s* and 1 + 2*s* for the ancestral homozygote, heterozygote and derived homozygote genotypes, respectively.

### Expected SFS

From PRF theory (Sawyer and Hartl 1992; Sethupathy and Hannenhalli 2008),

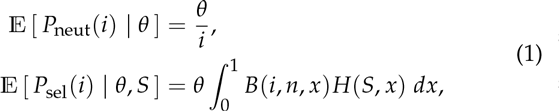

where

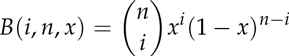

is the binomial probability of observing *i* derived alleles in a sample of size *n*, when the true allele frequency is *x*, and

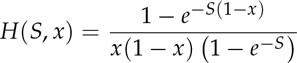

Note that due to our scaling of the mutation rate, *H*(*s*, *x*) is pro-ortional (with a factor of 1/2) to the mean time a new semidomnant mutation of scaled selection coefficient *S* = 4*N*_*e*_*s* spends between *x* and *x* + *dx* (Wright 1938). Figure 1D shows the expectations from equation (1) as a function of *S*.

To obtain 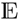 [*P*_sel_(*i*) | *θ*, *ϕ*], we integrate over the DFE,

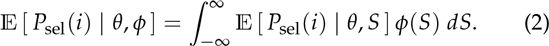

Relative to the expected SFS of independent sites under a Wright-Fisher constant population (equations (1) and (2)), the observed SFS can be distorted due to demography, ascertainment bias, non-random sampling, and linkage. We account for such distortions that affect both the neutral and selected sites to a similar extent by using the approach of Eyre-Walker *et al.* (2006) and introduce nuisance parameters *r*_*i*_, 1 ≤ *i* < *n*, that scale the expected SFS, for *z* ∈ {neut, sel},

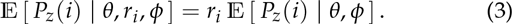

To avoid identifiability issues, we set *r*_1_ = 1.

### Full DFE and divergence counts

Unlike methods that infer only a strictly deleterious DFE, we can incorporate a full DFE that includes both deleterious and beneficial mutations. Additionally, to add flexibility in the inference of the full DFE, we optionally model divergence counts (the number of observed fixed mutations relative to an outgroup) *d*_*z*_ as a Poisson distribution with mean 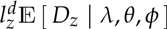. Here, 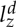 is the number of sites used for divergence counts, and can possibly be different than *l*_*z*_. We have that

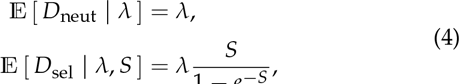

where λ = *Tθ* is a composite divergence parameter that accounts for the number of neutral mutations that go to fixation during the divergence time *T* from the MRCA of the ingroup population to the outgroup (blue full line in Figure 1A). The term *S*/(1 - *e*^−*S*^) accounts for the fixation of a mutation with scaled selection coefficient *S*, and can be obtained as lim_x→1_ *H*(*S*, *x*). Figure 1D shows the expectations for the divergence counts at selected sites from equation (4) as a function of *S*.

As divergence counts are calculated by comparing the outgroup sequence to the sample of sequences from the ingroup, polymorphism may be misattributed as divergence, i.e. mutations that are polymorphic in the ingroup population but fixed in the sample are counted as divergence. This is the case for mutations that occur between the MRCAs of the sample and ingroup (blue dot-dash line in Figure 1A). As noted by Keightley and Eyre-Walker (2012), misattributed polymorphism can lead to biased inference of *α*. To account for this, we adjust the above means to also incorporate the misattributed polymorphism by increasing the expectations with the contributions coming from mutations present in all *n* sampled individuals,

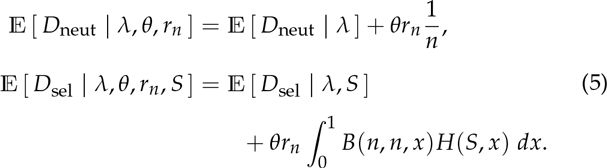

Assuming that the ingroup and outgroup share the same DFE, we integrate over it to obtain

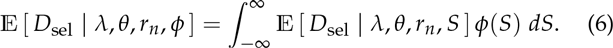

### Unfolded SFS and ancestral misidentification

When only a deleterious DFE is inferred, the folded SFS is typically used, where only sums of the form *p*_*z*_ (*i*) + *p*_*z*_ (*n* - *i*) are modeled. This is sufficient for inference of deleterious DFE (Keightley and Eyre-Walker 2007). However, the unfolded SFS contains valuable information for inference of the full DFE, as beneficial mutations are expected to be present in high frequencies (Durrett 2008; Fay and Wu 2000). To obtain an unfolded SFS, the ancestral state needs to be identified, and this is error prone. To account for potential misidentification of the ancestral state, we model the mean of *P*_*z*_(*i*), *z* ∈ {neut, sel}, as a mixture of sites whose ancestral states were correctly identified (with probability 1 - *ϵ*), or misidentified (with probability *ϵ*) (Williamson *et al.* 2005; Boyko *et al.* 2008; Glémin *et al.* 2015),

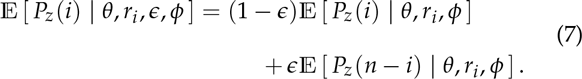

### Mutation variability

There is substantial evidence that both substitution and mutation rates vary along the genome (Golding 1983; Yang 1996; Francioli et al. 2015; Hodgkinson and Eyre-Walker 2011; Arndt et al. 2005), with a long tradition of modeling this variability in phylogenetic inferences as a gamma distribution (Golding 1983; Yang 1996). A few DFE inference methods allow for mutation rates to vary in a non-parametric fashion (Bustamante et al. 2003; Gronau et al. 2013). Here, we model mutation variability by assuming that mutation rates follow a gamma distribution with mean *θ*̅ and shape *a*. This is motivated by the phylogenetic approaches, but also by mathematical convenience: if the mean of a Poisson distribution follows a gamma distribution, the resulting distribution is a negative binomial distribution. We assume that the data is divided into *m* non-overlapping fragments of lengths 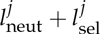, 1 ≤ *j* ≤ *m*, and for each fragment *j*, we have the SFS 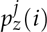, 1 ≤ *j* < *n*. Possibly, we have an additional *m*^*d*^ fragments of lengths 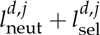 for which we have the divergence counts, 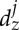. We assume that each fragment has a constant mutation rate *θ8*, but that mutation rates can vary between fragments. Given the mutation rate θ_*j*_ of the fragment *j*, then 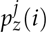 and 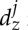 follow the Poisson distributions with means 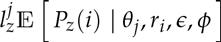 and 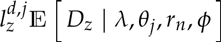, given by equations (1)–(6). Integrating over the mutation rates distribution, we obtain that 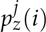 and 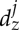 have a negative binomial distribution with shape *a* and means 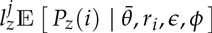 and 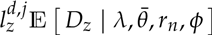, respectively.

### Inferring *α* using divergence or polymorphism data alone

Once the DFE is estimated, *α* can be calculated from the observed divergence counts as follows (Eyre-Walker and Keightley 2009)

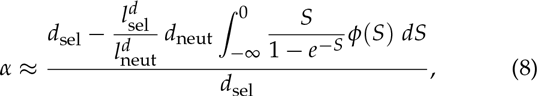

where the nominator represents the estimated number of adaptive substitutions, which are obtained by subtracting the expected deleterious and neutral substitutions from the total observed divergence counts at selected sites. Keightley and Eyre-Walker (2012) extended the above estimation of *α* to account for the misattributed polymorphism. Using our framework, we correct for the misattributed polymorphism by removing from *d*_sel_ and *d*_neut_ the expected number of mutations that are in fact polymorphic. These expectations can be readily obtained from equation (5) by setting λ = 0. Then the new estimate of *α* is obtained as in equation (8), where *d*_sel_ and *d*_neut_ are replaced with the re-adjusted divergence counts 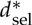 and 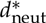 given by

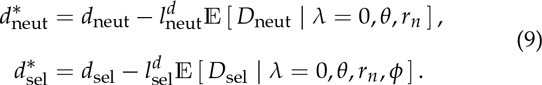

This approach to calculate *α* relies heavily on the assumption that the ingroup and outgroup share the same scaled DFE. However, if one has access to an estimated full DFE purely from polymorphism data, *α* can still be estimated by replacing the observed divergence counts with the expected counts from equation (4). As λ will cancel out in the resulting fraction, *α* can be obtained by setting λ = 1. Then,

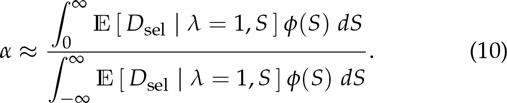

In the rest of this paper, we refer to the two above estimates of *α* as *α*_div_ and *α*_dfe_, respectively, to distinguish more clearly the type of information used.

### Likelihood estimation and comparison of models

The hierarchical framework described above allows maximum likelihood estimation of both evolutionary (mutation rates, DFE parameters) and nuisance parameters, as well as the error in the ancestral states, *ϵ*. Details about the implementation and optimization of the likelihood function are given in the Supplemental Material. Note that in our implementation, likelihood ratio tests (LRTs) can be used to test rigorously whether the polymorphism data provides evidence for a full DFE, or if a strictly deleterious DFE is sufficient for accounting for the data. This framework also allows to decide whether including nuisance parameters and / or ancestral errors provides a better fit to the data. The p-values for the LRT are obtained by assuming that likelihood ratio under the null hypothesis (reduced model is correct) is distributed as *χ*^2^.

## Results and Discussion

To investigate the statistical performance of our method to infer the DFE, *α* and test hypothesis regarding the contribution of beneficial mutations to patterns of polymorphism, we performed extensive simulations using SFS_CODE (Hernandez 2008). We explored a wide range of simulated DFEs (12 full DFEs and 5 deleterious DFEs, Table S1), chosen such that the simulated *α* had one of four possible values (0%, 20%, 50% and 80%, Figure 2A). Most simulations were performed using a constant population size and without error in the identification of the ancestral state. Results are shown for this type of data if not otherwise specified. These assumptions were later relaxed. The simulations contained linkage and were performed to resemble exome data. For each considered simulation scenario (one given DFE, demographic, linkage, misidentification of the ancestral state), we simulated 100 replicate data sets. For more details on the simulated data, see the Supplemental Material.

**Figure 2.**
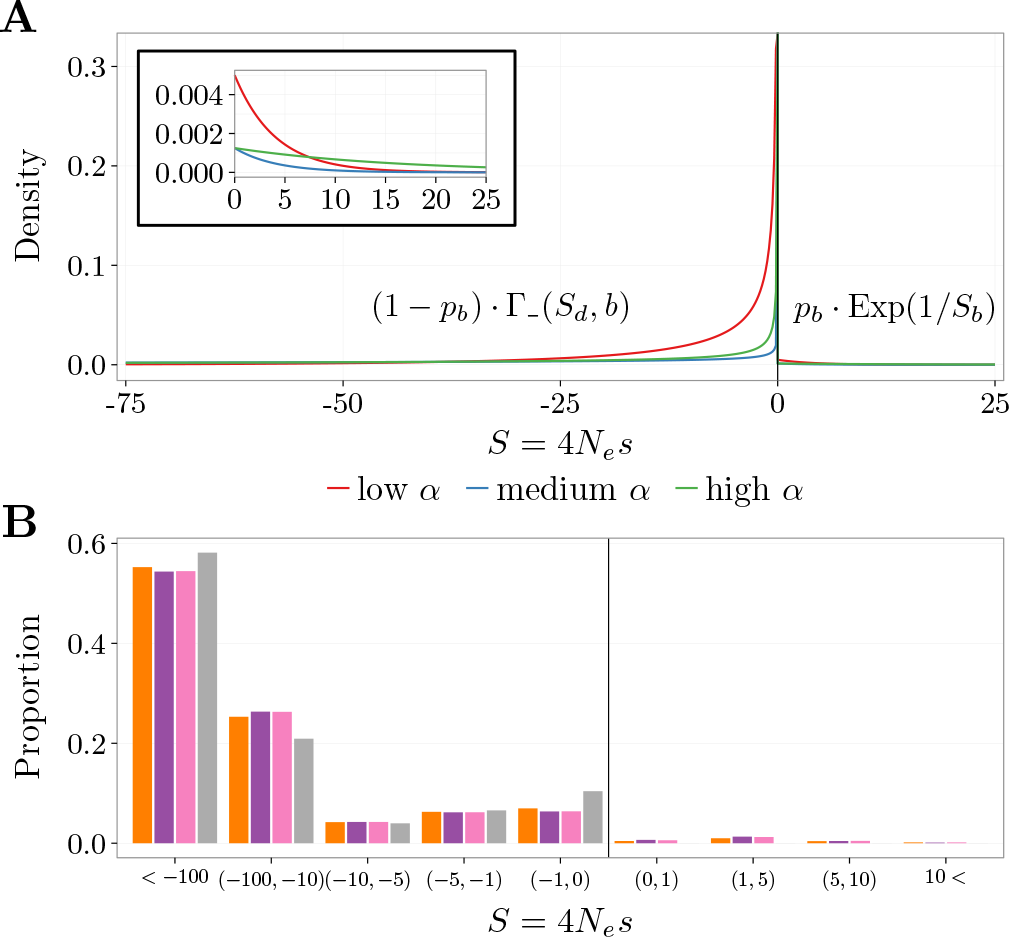
Example of simulated and inferred gamma + exponential DFEs. (A) Three of the simulated DFEs (corresponding to LALSD, MALPB, and HAHSB from Table S1) with different as (proportion of beneficial substitutions). The DFEs are parameterized by *S*_*d*_ (mean selection coefficient of deleterious mutations), *b* (shape of distribution of deleterious DFE), *p*_b_ (proportion of beneficial mutations), and *S*_*b*_ (mean selection coefficient of beneficial mutations). The inset shows a zoom-in of the beneficial part of the DFE. (B) Simulated discretized DFE (orange, corresponding to MAMSD from Table S1), together with the mean (over the 100 replicates) inferred discretized DFE using both polymorphism and divergence data (purple) and only polymorphism data (pink and gray), where a full DFE (pink) and a deleterious DFE (gray) was inferred.

The general shape of the DFE is not agreed upon (Welch *et al.* 2008; Bataillon and Bailey 2014). The DFE has been modeled using a wide range of functional continuous forms (Boyko et al. 2008; Kousathanas and Keightley 2013; Galtier 2016), but also as a discrete distribution (Gronau et al. 2013; Kousathanas and Keightley 2013; Keightley and Eyre-Walker 2010). Here, we use a DFE consisting of a mixture between gamma and exponential distributions, that model deleterious and beneficial mutations, respectively. With probability 1 - *p*_*b*_, a new mutation is deleterious and its selection coefficient comes from a reflected gamma distribution with mean *S*_*d*_ < 0 and shape *b*, while with probability *p*_b_, a new mutation is beneficial and its selection coefficient comes from an exponential distribution with mean *S*_*b*_ > 0 (Figure 2A). We do not explore alternative parametric DFE families. For such studies, we refer the reader to Kousathanas and Keightley (2013); Welch et al. (2008).

We inferred the DFE and alpha parameters in our model using three different models: a full DFE was inferred from both polymorphism and divergence data; a full DFE was inferred from polymorphism data alone; an only deleterious DFE was inferred from polymorphism data alone. From the inferred DFEs, we calculated *α*_dfe_ and *α*_div_. For the inference assuming only a deleterious DFE, *α*_dfe_ is always 0, and for such inference we therefore only calculated *α*_div_. The distortion parameters *r* were always estimated, while the ancestral misidentification error *ϵ* was fixed to 0, unless otherwise specified. We report the inference performance using log2(estim/sim) on a log-modulus scale. Here, estim is the estimated value, while sim is the simulated value. Unlike the relative error, this log ratio gives equal weight to both overestimation and underestimation of the parameters. For example, the log ratios of 1 and -1 correspond to the estimated value being double or half the simulated value, respectively. When sim = estim, the ratio is equal to 0. See the Supplemental Material for details.

### Inference of deleterious DFE

Using simulations that did not contain any beneficial mutations in the polymorphism data, we first investigated how well we can infer the deleterious DFE and if our method can recover the fact that all polymorphic mutations are deleterious. We observed that the two parameters determining the deleterious DFE, *S*_*d*_ and *b*, and *α*, are inferred accurately when only a deleterious DFE was estimated (Figure 3A and Figure S1). When, instead, a full DFE was inferred from the polymorphism data alone, the parameters showed different amounts of bias (Figure 3A and Figure S1). A crucial question is whether the data allows one to decide correctly which model is most sensible: the full DFE or only deleterious DFE? When using a LRT to compare the relative goodness of fit on simulated data, virtually all data sets tended to reject the full DFE model in favor of the reduced model featuring only deleterious mutations in the DFE (Figure S2). This indicates that while our method can account for the presence of beneficial mutations in the SFS data, it can also accurately detect if there is no empirical evidence for such mutations in the data. So in principle, one can perform estimation under both the full and deleterious DFE models and use the LRT to decide which model is most appropriate for the data.

### Inference of full DFE

From the expected contribution of mutations to polymorphism and divergence data, as a function of *S* (Figure 1D), it is evident that if beneficial mutations occur at any appreciable rate, they should have a non-negligible impact in the polymorphism data. This suggests that it should be possible to infer the full DFE from polymorphism data alone. We investigated this using data generated under a full DFE. As one might expect, the deleterious DFE parameters were inferred equally well regardless of whether the divergence data was used or not (Figure S3). The variance of the estimates seems to be somewhat larger when divergence data is not used, but this is most likely due to the inference using less data. The parameters of the beneficial part of the DFE and *α* were inferred with different levels of accuracy (Figure 3B and Figure S3). From the simulation scenarios considered here, it is apparent that the value of *α* predicts the accuracy: the higher *α*, the better the prediction, for both inference with and without divergence. In short, when beneficial mutations are comparatively rare and of very small effects, estimating their properties and *α* is challenging (even with divergence data). Conversely, when beneficial mutations are relatively common, they dominate the divergence counts, but also make substantial contribution to the SFS counts, which alone can allow reliable estimation of the beneficial fraction of the DFE and *α*. For the lower values of *α* (*α* ≈ 20%), the use of divergence data provides more accurate estimates than when relying on polymorphism data alone. This could perhaps be explained by the fact that, in this case, the polymorphism data is heavily dominated by deleterious mutations and it is more difficult to tell apart the amount of beneficial selection from polymorphism data alone. However, as *α* increases, the differences in performance between the inference with and without divergence diminishes, strongly indicating that divergence data is not necessarily needed for accurate inference.

**Figure 3.**
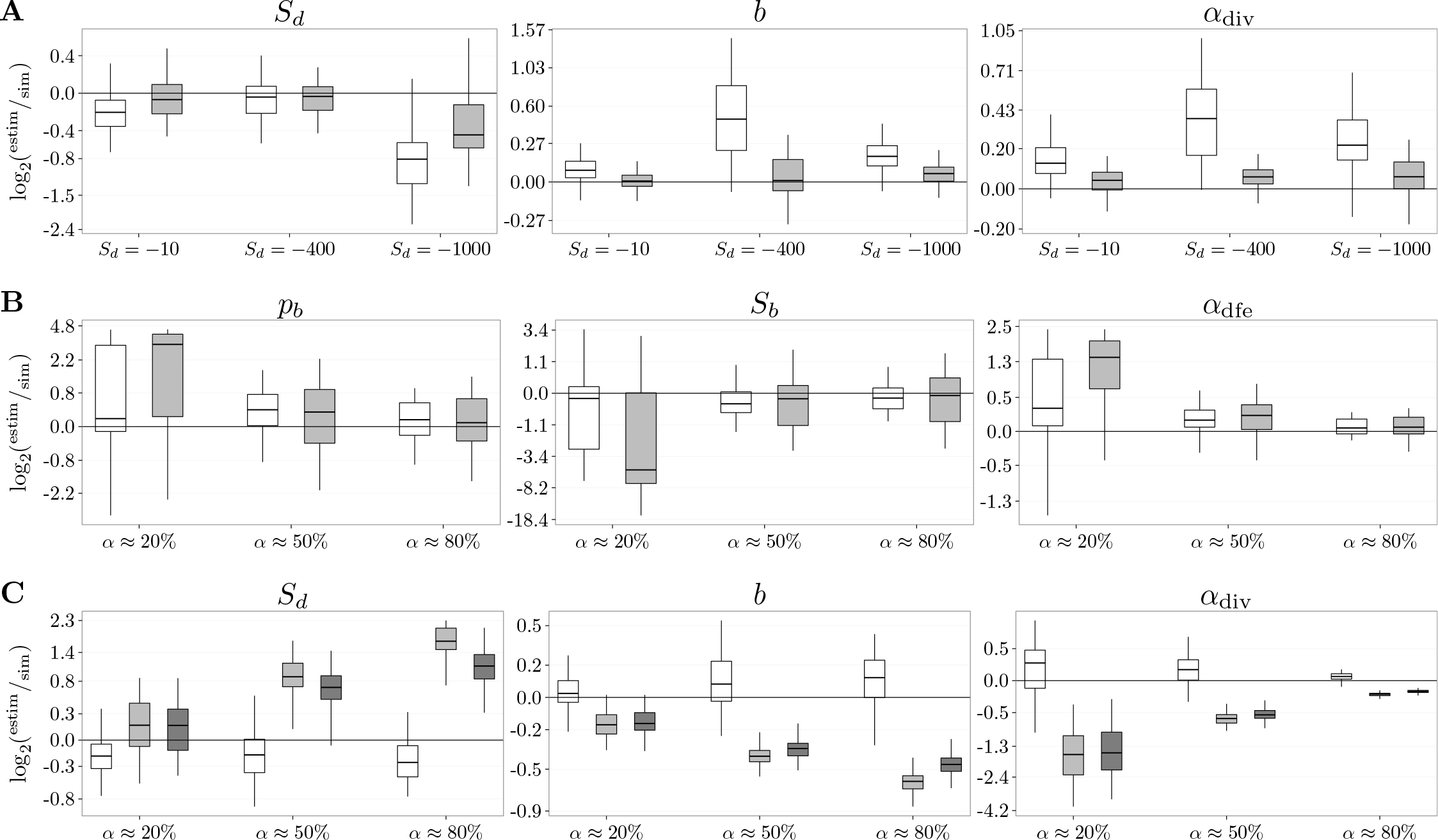
Inference of *α* (proportion of beneficial substitutions) and DFE parameters: *S*_*d*_ (mean selection coefficient of deleterious mutations), *b* (shape of distribution of deleterious DFE), *p*_*b*_ (proportion of beneficial mutations), and *S*_*b*_ (mean selection coefficient of beneficial mutations). (A) Quality for inference performed on polymorphism data alone, for three simulated deleterious (corresponding to DelLSD, DelMSD, and DelHSD from Table S1) DFEs with different Sds. The DFE parameters are inferred using only polymorphism data assuming a full (white boxes) and deleterious (gray boxes) DFE. (B) Quality for inference performed on polymorphism and divergence data, for three simulated DFEs with different αs (corresponding to LALSD, MAMSD, and HAHSD from Table S1). The DFEs differ only in the simulated value of *S*_*d*_. The DFE parameters are inferred using both polymorphism and divergence (white boxes) and only polymorphism (gray boxes) data. (C) Quality for inference performed on polymorphism data alone, for three simulated DFEs with different αs (corresponding to LALSB, MAMSD, and HAHSB from Table S1). The DFEs differ only in the simulated value of *S*_*b*_. Only polymorphism data is used, and the DFE parameters are inferred assuming a full DFE where e is set to 0 and is not estimated (white boxes) and a deleterious DFE (gray boxes), where *ϵ* is set to 0 and is not estimated (light gray boxes), or is estimated (dark gray boxes). The data was simulated with *ϵ* = 0.

Similar to Schneider *et al.* (2011), we observe a strong negative correlation between the proportion of beneficial mutations *p*_*b*_ and their scaled selection coefficient *S*_*b*_ (Figure S4). This illustrates the fact that *p*_*b*_ and *S*_b_ are difficult to estimate separately, but their product, which largely determines *α*, is more accurately estimated. This can be seen in Figure 3B and Figure S3, where even though *p*_*b*_ and *S*_*b*_ might be biased, overall *α* is inferred more accurately. Schneider *et al.* (2011) reported that the estimation of *p*_*b*_ and *S*_*b*_ improves as more sites are available. In our data simulations we used a fixed number of sites, but we do observed that *p*_*b*_, *S*_*b*_ and *α* are better estimated as *α* increases.

When inferring a full DFE, we can calculate both *α*_div_ and *α*_dfe_, which should both be good predictors of the true simulated *α*. We generally found very good correlation between the two estimated values (Figure S5), and perhaps not surprising, the estimates of *α*_dfe_ obtained when performing inference on both SFS and divergence data were more tightly correlated.

We note here that Schneider *et al.* (2011) is the only method that we are aware of that can estimate both a full DFE and *α* from polymorphism data alone, though the authors did not investigate the power to infer *α*, but rather the product of *p*_*b*_*S*_*b*_, which is taken as a proxy for *α*. Additionally, they did not consider in their simulations different deleterious DFEs. Our simulated DFEs where chosen such that they cover cases with the same simulated *p*_*b*_ and *S*_*b*_, but generate different *α*s. The differences in *α* can be driven by the amount of beneficial mutations, but also by the intensity of purifying selection, or, said slightly differently, the properties of the deleterious fraction of the DFE. These simulated data set revealed that the amount and strength of positive selection is not the only determinant in how accurately *p*_*b*_ and *S*_*b*_ are inferred. For example, the results in Figure 3B are given for simulated DFEs that differ only in the value of *S*_*d*_, i.e. the strength of purifying selection, and we find in this instance a clear difference in the inference performance in terms of relative error.

### Bias from not inferring full DFE

Given that divergence data is clearly not necessary for reliable estimates of the full DFE, a question arises: what happens when inference methods ignore the presence of beneficial mutations in the polymorphism data? For this, using the simulated data sets generated using full DFEs, we performed inference only on polymorphism data where we inferred either a full DFE, like before, or under a reduced model restricted to only a deleterious DFE (Figure 3C and Figure S6). Note that this corresponds to the current state of the art for empirical studies of DFE from population genomics data, where data tend invariably to be analyzed under the assumption that SFS data is to be fitted exclusively with a deleterious DFE (Racimo and Schraiber 2014; Bataillon *et al.* 2015; Halligan *et al.* 2013; Arunkumar *et al.* 2015; Charlesworth 2015; Harris and Nielsen 2016; Slotte *et al.* 2010; Strasburg *et al.* 2011). When *α* was ≈ 20%, the inferred *S*_*d*_ and *b* were, at times, more accurate when only a deleterious DFE was used. However, as *α* increased, the two parameters were increasingly biased. The mean *S*_*d*_ was estimated to be more negative, while the shape *b* was estimated to be closer to 0: the inferred deleterious DFEs were getting much more leptokurtic than the parametric DFE used to simulate the data. This resulted in inferring DFEs with more probability mass accumulating close to 0. A straightforward interpretation is that the inference method attempted to fit the SFS counts contributed by the weakly beneficial mutations by fitting a DFE that comprised a sizable amount of weakly deleterious mutations (the best proxy for beneficial mutations). A comparison of the simulated and inferred discretized DFEs (Figure 2B and Figure S7) illustrates this point: the inference with only deleterious DFE overestimated appreciably the amount of mutations with a selection coefficient in the range ( - 1,0) (simulated: 0.07, deleterious DFE: 0.11) and (-5, -1) (simulated: 0. 063, deleterious DFE: 0.067) ranges.

DFE methods that do not model beneficial mutations in the polymorphism data use a folded SFS. To mimic this behavior, we allowed for *ϵ* to be estimated, even though no errors in the identification of ancestral state were simulated. We observed that *ϵ* reduces partially the bias in the parameters (Figure 3C, Figures S6 and S7).

Using a LRT we could test, as before, for the presence of beneficial mutations in the polymorphism data by comparing the inferences with a full or deleterious DFE (Figure S8). We observed that the larger *α*, the stronger the preference for the full DFE model. We also noticed that, even though *α* might be relatively large, if the mean strength of beneficial selection was very low (Figure S8, MALSB where *S*_*b*_ = 0.1), the LRT indicated that there were no beneficial mutations in the polymorphism data. Such mutations can pass as weakly deleterious mutations when fitting the data. The LRT also showed an increasing preferencewith *α* for *ϵ* when only deleterious DFE is inferred, indicating, as expected, that the model with *ϵ* ≠ 0 could account for some of the weakly beneficial mutations present in the polymorphism data.

Inferring only a deleterious DFE leads to a consistent bias in *α* as well. This bias is not that well correlated with the simulated 539 value of *α*, but it is apparent that a higher *α* leads to a smaller bias (Figure S6). This is in contrast to the bias observed for *S*_*d*_ and *b*. To obtain *α* from a deleterious DFE only, we need to rely on the divergence data. Perhaps, when *α* is large, the signal of positive selection is so strong in the divergence data that it overrides, to some extent, the bias in *S*_*d*_ and *b*, leading to a more accurate estimate of *α*.

The assumption of negligible contribution of beneficial mutations to SFS counts can be traced back to Smith and Eyre-Walker (2002). To support the claim, the authors stated that “if advantageous mutations, with an advantage of *N*_*e*_*s* = 25 occur at one-hundredth the rate of neutral mutations, they will account for 50% of substitutions, but account for just 2% of heterozygosity”. Our simulated *S*_*b*_ (which is scaled by 4*N*_*e*_) was typically 4. To investigate what happens when selection is much stronger, we simulated a full DFE with *S*_*b*_ = 800 such that only 10% of beneficial mutations (0.2% of all mutations) had a selection coefficient of 100 or less. For this, the simulated *α* was nearly 100% and one would expect that, as selection is so strong, most mutations would fix quickly. While the DFE parameters could not be recovered as accurately (Figure S9), the estimated *α* was very precise, regardless of the model used for inference. This points to the fact that, even when positive selection is very strong, there is enough information left in the polymorphism data to be able to estimate *α* without relying on divergence data. The bias in *S*_*d*_, *b* and *α* (Figure S9) and LRT (Figure S10) from inference with only deleterious DFE followed the same trend as before. However, even though *α* was large, *p*_*b*_ and *S*_*b*_ were not that well estimated. This is most likely because when *S*_*b*_ is getting very large, the expected counts from equation (1) become independent of *S*, since 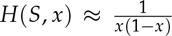 for large *S* (Figure 1D). This explains why the inference method will have trouble narrowing precisely the value of *S*_*b*_.

Keightley and Eyre-Walker (2010) investigated if the presence of beneficial mutations in the polymorphism data could potentially affect the inference when assuming only a deleterious DFE. For this, they simulated data using a partially reflected gamma distribution, given by

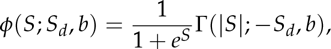

where Γ(*x*; *m*, *s*) is the density of a gamma distribution with mean *m* and shape *s*. This distribution arises from the assumption that the absolute strength of selection is gamma distributed and that each site can be occupied by either an advantageous or a deleterious allele, both having the same absolute selection strength |*s*|. Keightley and Eyre-Walker (2010) simulated datawith |*S*_*d*_| = 400 and *b* = 0.5. Due to the chosen distribution, the simulated proportion of beneficial selection was *p*_*b*_ = 0.0214, while the mean selection coefficient of beneficial mutations was only *S*_*b*_ = 0.014 (*p*_b_*S*_*b*_ = 0.00029). These values are close to one of our own simulated DFEs with *α* ≈ 20% (LALSB, Table S1), with the difference that we simulated an *S*_*b*_ that was approximately 7 times larger. For this simulation scenario we did, indeed, find little bias in *S*_*d*_ and *b* when inferring only a deleterious DFE (Figures S6 and S7). However, arguably, this strength of beneficial mutations is extremely low. For example, Schneider *et al.* (2011) inferred the strength of beneficial mutations from *Drosophila* and found *P*_*b*_*S*_*b*_ to be two-three orders of magnitude higher: *p*_*b*_ = 0.0096 and *S*_*b*_ = 18 (*p*_*b*_*S*_*b*_ = 0.1728).

### Impact of ancestral error on inference

The results presented above were based on simulations where the true ancestral state was used. To investigate the consequences of misidentification of the ancestral state, we added errors to the simulated data (see Supplemental Material for details). Inferring a full DFE and using divergence data, we found that we can properly account for the rate of misidentification, and the error *ϵ* is accurately recovered (Figure 4 and Figure S11). As expected, the inference of the DFE and *α* is biased when the misidentification is not accounted for. A LRT for *ϵ* ≠ 0 (Figure S14) supported the use of a model including the joint estimation of *ϵ* and DFE parameters for the data with errors, but rejected the more complex model for the data without error.

**Figure 4.**
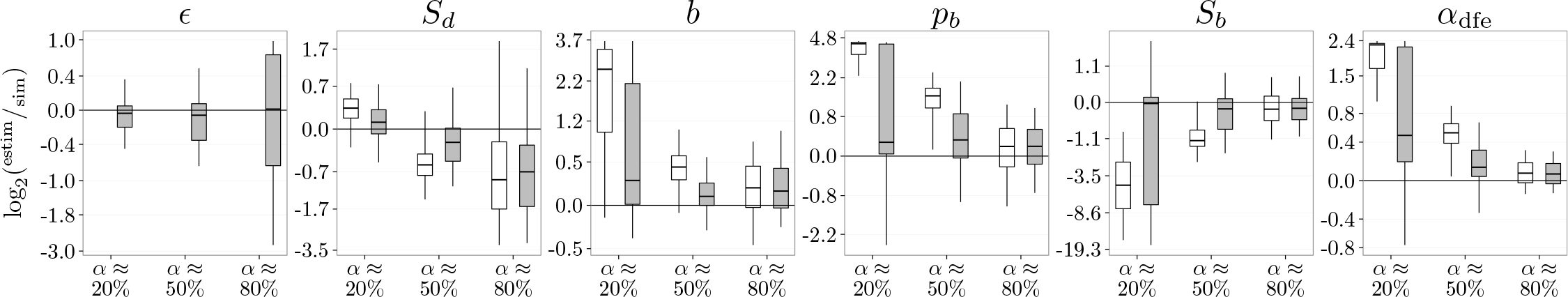
Inference of *α* (proportion of beneficial substitutions), *ϵ* (rate of ancestral error), and DFE parameters: *S*_*d*_ (mean selection coefficient of deleterious mutations), *b* (shape of distribution of deleterious DFE), *p*_*b*_ (proportion of beneficial mutations), and *S*_*b*_ (mean selection coefficient of beneficial mutations). The figure shows the inference quality for three simulated DFEs (corresponding to LALSD, MAMSD, and HAHSD from Table S1) with different *αs*. The DFEs differ only in the simulated value of *S*_*d*_. A full DFE is inferred from both polymorphism and divergence data, and *ϵ* is set to 0 and is not estimated (white boxes), or is estimated (gray boxes). The data was simulated with *ϵ* = 0.05.

Galtier (2016), who also used distortion parameters *r*_*i*_ when inferring the DFE, stated that these parameters are “expected to capture any departure from the expected SFS as soon as it is shared by synonymous and non-synonymous sites”. Our results indicate that the misidentification of the ancestral state cannot be accurately accounted for by the *r*_*i*_ parameters (Figure 4 and Figure S11). However, we did find that the resulting bias decreased with *α* and that the preference (as measured by a LRT) for models inferring *ϵ* ≠ 0 also decreased for data sets with higher *α* (Figure S14). For data simulations with *α* ≈ 80%, the inference was just as good when *ϵ* was set to 0. To investigate this in more details, we also ran the inference with *r*_*i*_ = 1 (i.e. no distortion correction) and *ϵ* = 0 on those simulated DFEs. The results showed a large bias in the DFE parameters and *α* when *r* = 1 (Figure S12), and a LRT favored the estimation of *r*_*i*_s (Figure S15). This illustrated that merely using the *r*_*i*_ parameters without explicitly accounting for misidentification of the ancestral state is not always accurate and can bias inference of DFE and *α*.

Both the presence of beneficial mutations and *ϵ* ≠ 0 create similar patterns in the polymorphism data: the frequency of the common derived alleles increases. We have seen before that *ϵ* can account for some of the positive selection in the data (Figure 3C, Figures S6 and S7). Similarly, we observed that positive selection can account for some of the misidentification of ancestral state. On simulations with a deleterious DFE and incorrect ancestral states, we found that when assuming *ϵ* = 0, the parameters inferred when a full DFE is assumed are, generally, more accurate than when only a deleterious DFE is inferred (Figure S13). A LRT also supported the use of a full DFE (Figure S16). Comparing the inferred *p*_*b*_ and *S*_*b*_ when *ϵ* is inferred or not (Figure S13) showed that these parameters are higher when *ϵ* = 0, further indicating that they captured some of the ancestral misidentification errors. Therefore, if the data contains sites that have the ancestral state misidentified, which is virtually always the case in empirical data sets, ancestral misidentification will be wrongly interpreted as positive selection if the misidentification is not accounted for. If *ϵ* is inferred jointly with the DFE parameters, a LRT comparing models with full DFE or only deleterious DFE can correctly detect that the polymorphism data does not contain any beneficial mutations (Figure S16). Our simulation results illustrated that systematically incorporating the rate of ancestral error is crucial for a reliable inference of DFE parameters and *α*.

### Distortions of the SFS by linkage and demography

It has previously been suggested that correcting for the effect of demography–using the observed SFS at neutral sites–can also reduce some of the bias introduced by linkage in the data (Kousathanas and Keightley 2013; Messer and Petrov 2012). We explored this possibility by simulating different levels of linkage (see Supplemental Material for details). We found that, indeed the *r*_*i*_ parameters could partially correct for the presence of link age (Figure S17), with the most pronounced effect on *S*_d_ and *b*. A LRT for *r* ≠ 1 (Figure S18) increasingly favored the more models fitting *r*_*i*_ as the level of linkage increased.

**Figure 5.**
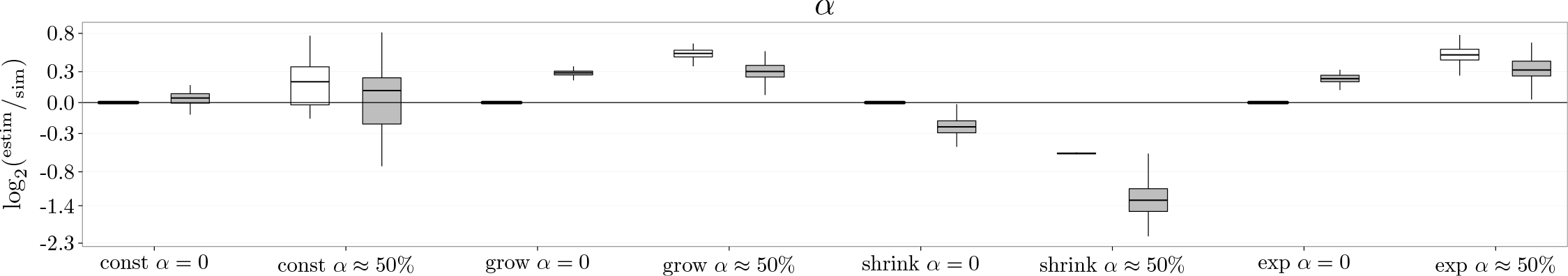
Inference of *α* (proportion of beneficial substitutions). The figure shows the inference quality for *α*_dfe_ (inferred from DFE alone, white) and adiv (inferred from DFE and divergence data, gray) for four simulated demographic scenarios (detailed in the Supplemental Information) and a deleterious DFE only (*α* = 0, corresponding to DelMSD from Table S1) or a full DFE (*α* ≈ 50%, corresponding to MAMSD from Table S1). For all inference only SFS data was used, and a LRT was performed to compare the full and deleterious only DFE models. The estimated value of *α* was chosen from the model preferred by the LRT.

For the previous simulations we used a constant population size. To check that the *r*_*i*_ parameters can also correct for demography, we simulated additional data, using different demography scenarios (see Supplemental Material for details). When populations size varies in time, *N*_*e*_ is typically taken to be the harmonic mean of the different sizes (Kliman *et al.* 2008). While this might be a good approximation for the neutral sites, the site under selection experience a different *N*_*e*_, which depends on the strength of selection *S* (Otto and Whitlock 1997). Therefore, foi these simulations, we do not have a priori knowledge of the *N*_*e*_ that accurately captures the interaction between selection and demography, and we could only compare parameters that are in dependent of *N*_*e*_ (*b*, *p*_*b*_), and *α* for which a value can be obtained by tracking the proportion of adaptive mutations contributing to divergence in the forward simulations used to generate the data sets. Like before, we first simulated a deleterious DFE, similar to previous studies (Boyko *et al.* 2008; Eyre-Walker *et al.* 2006; Eyre-Walker and Keightley 2009; Keightley and Eyre-Walker 2007) We found that a LRT correctly detected that *r*_i_≠1 (Figure S20) but that the parameters can correct only partially for the effect of demography (Figure S19). The *b* parameter was inferred ac curately when the *r*_*i*_ parameters were estimated. However, the estimated *α* was still biased (Figure 5). As no full DFE was inferred, *α* was calculated from the divergence data. For this, the same DFE was assumed in the ingroup and outgroup. However, as the ingroup now underwent variable population size, its Ne was different from the Ne of the outgroup (which had a constant size), and therefore the two populations had different scaled DFEs. This difference could explain the observed bias in *α* Eyre-Walker and Keightley (2009) noticed the same effect and proposed a correction for *α*. However, their correction requires the ratio of the Nes of the two populations, which is typically not known.

We then investigated if the *r*_*i*_ parameters could also correct for demography when a full DFE was simulated (Figure S21). The LRT (Figure S22) showed a clear preference for *r*_*i*_ ≠ 1. When inferring the DFE from both polymorphism and divergence data, we observed a bias in *b* and *p*_*b*_. As before, this was caused by the incorrect assumption of a shared DFE between the ingroup and outgroup. When only polymorphism data was used for the inference, the *r*_*i*_ parameters could accurately correct the estimation of *p*_*b*_ and *b*, but the inferred *α* (estimated via *α*_dfe_) was still slightly biased. To investigate if this bias was caused by linkage, we also ran simulations with reduced linkage (Figure S21), but the bias remained.

We investigated if the full or deleterious DFE model is preferred for the data simulated under variable population size. We found that a LRT consistently preferred the full DFE model when the SFS data contained beneficial mutations (Figure S23). Under demographic simulations, the estimated *α*_dfe_ and *α*_div_ could differ considerably (Figure 5). When only a deleterious DFE was simulated, relying on divergence data to estimate *α* can lead to heavily biased estimates. Note that when the population size was halved and a full DFE was simulated, the LRT favored the incorrect deleterious DFE model. This indicates that the *r*_*i*_ parameter cannot fully capture the demography and in this particular simulation, the incorrect model choice can be explained by the extra deleterious load incurred by the population shrinkage.

The simulated demographics were the same for both deleterious and full DFE simulations, and therefore the inferred *r*_*i*_ parameters on the deleterious and full DFE data should be highly correlated. We did, in fact, detect a strong correlation (Figure S24). One of the simulations showed no correlation in *r*_*i*_ and the LRT preferred the less complex model with *r*_*i*_ = 1 (Figures S20 and S22, SHRINK). The change in population size for this simulation was most likely not strong enough for it to leave an appreciable footprint in the data.

Galtier (2016) is the only study that we are aware of that tested if demography can accurately be accounted for when a full DFE was simulated. While the estimated *α*s from Galtier (2016) were somewhat more accurate than what we found, there are critical differences between these studies. While we simulated a continuous full DFE, Galtier (2016) assumed that all beneficial mutations had the same selection coefficient *S*_*b*_. Nevertheless, Galtier (2016) inferred a continuous full DFE and used equation (8) for calculating *α*, where the integration limit was set to some *S*_adv_ > 0 instead of 0. The reasoning behind this is that mutations with a selection coefficient *S* > 0 that is not very large should not be considered advantageous mutations. Galtier (2016) used an arbitrary cutoff at *S*_adv_ = 5. Note that a different cut-off value of *S*_adv_ would lead to different *α*s: the smaller *S*_adv_, the larger the estimated *α*.

### Comparison with the dfe-alpha method

We chose to compare our method with dfe-alpha, one of the most widely used inference methods for DFE and *α*. dfe-alpha was originally developed to infer a deleterious DFE (Keightley and Eyre-Walker 2007), and it was subsequently extended to estimate *α* (Eyre-Walker and Keightley 2009), model a full DFE (Schneider *et al.* 2011) and correct *α* for misattributed polymorphism (Keightley and Eyre-Walker 2012). While dfe-alpha can infer a full discrete DFE, the method to account for potential errors in the ancestral state described in Schneider *et al.* (2011) is not implemented in dfe-alpha. At the time when we ran out comparison, we could not find any option in dfe-alpha for accounting for such errors. As we showed that this is crucial for accurate inference (Figure 4 and Figure S11), we therefore chose to run dfe-alpha with a folded SFS, where only a deleterious DFE can be estimated. We then compared with our method when only a deleterious DFE was inferred, where, as before, *03F5;* was either set to 0, or estimated. Although these comparisons are therefore quite limited in scope, we found that, for simulations with only a deleterious DFE, our method provided better estimates and with lower variance than dfe-alpha (Figure 6 and Figure S25). For these simulations, we also found that, sometimes, dfe-alpha estimated an *α* that was very large, both on the negative and positive side (Figure S25, DelHB simulation). This seemed to be the result of the correction for the misattributed polymorphism, as the uncorrected *α* was much closer to the true value (data not shown). This most likely explains the general differences observed between the estimated *α* from dfe-alpha and our method. When the inference was performed on data simulated with a full DFE, we observed the same type of bias in *S*_*d*_ and *b* as described before (Figure 6 and Figure S25). When demography and a strictly deleterious DFE were simulated, the estimation was, again, comparable (Figure S26). However, when demography was simulated on top of a full DFE, the bias of *b* differed between dfe-alpha and our method. This could, perhaps, be explained by the differences between the two methods for accounting for demography: while we used the nuisance parameters *r*_*i*_, dfe-alpha assumes a strict simplified demographic scenario and only allows the population to undergo one size change in the past.

**Figure 6.**
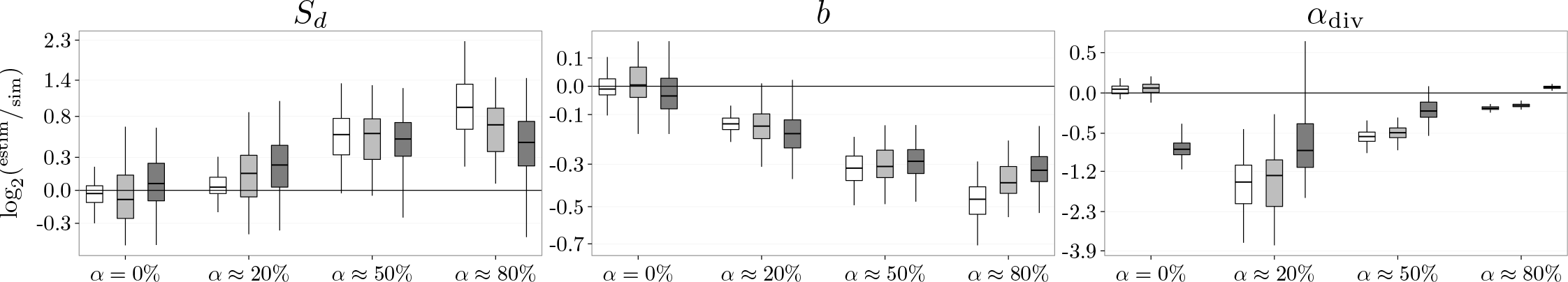
Comparison to dfe-alpha of inference of *α* (proportion of beneficial substitutions) and deleterious DFE parameters: *S*_*d*_ (mean selection coefficient of deleterious mutations) and *b* (shape of distribution of deleterious DFE), for four simulated deleterious DFEs (corresponding to DelMSD, LALSD, MAMSD and HAHSD from Table S1) with different *α*s. The DFE parameters are inferred using only polymorphism data, assuming a deleterious DFE, where *ϵ* is set to 0 and is not estimated (white boxes), or is estimated (light gray boxes). The inference from dfe-alpha is given in the dark gray boxes. The data was simulated with *ϵ* = 0.

## Conclusion

We have presented a new method to infer the DFE and proportion of advantageous substitutions, *α*, from polymorphism and divergence data. Using our framework, we demonstrated that inference can be performed using polymorphism data alone, and that this lead to more accurate inference when the DFE is not shared between the ingroup and the outgroup. We additionally illustrated that when the effects of beneficial mutations on polymorphism data were not modeled, the inferred deleterious DFE was biased. This bias comes from an increase of mutations at selected sites that segregate at high frequencies. Methods ignoring the contribution of beneficial fraction to SFS counts will tend to infer DFEs that have a larger amount of slightly deleterious mutations, as this is the best way to account for the observed data. Therefore, the estimated deleterious DFE had a much larger mass close to 0 compared to the simulated deleterious DFE. This, in turn, could be achieved by a larger (more negative) *S*_*d*_ and a lower *b* of the r distribution used here for the deleterious DFE. In cases where polymorphism data did not contain any beneficial mutations, the inference was much more accurate under a reduced model positing only a deleterious DFE. We showed that when applying our method, the use of a LRT comparing a model featuring a full DFE and a deleterious DFE, would accurately select the reduced model and allow precise inference of the deleterious DFE. This is an important result, as it suggests that using a full DFE for inference from SFS data does not come with a cost when no beneficial mutations contributed to the SFS counts, and that the method does not spuriously infer presence of beneficial mutations.

In order to correct for demography and other forces that can distort the SFS data, such as linkage, we used the so-called nuisance parameters *r*_*i*_s. These parameters have the potential of accounting for more complex scenarios without directly modeling the underlying changes in population size, and potentially, other events such as migration and admixture. This could prove more robust than just allowing for one (or two) population size changes, as dfe-alpha assumes. However, we did not test the behavior of our method under these more complex scenarios and the extent of bias in *α* they might generate.

In order to infer the full DFE, we used the unfolded site frequency spectrum (SFS). This requires the identification of the ancestral state, which is prone to errors. The errors in the identification of the ancestral state can, for example, be accounted for by using a probabilistic modeling of the ancestral state (Schneider *et al.* 2011; Gronau *et al.* 2013). We chose to assume that the polymorphism data is composed of a mixture of sites with correctly inferred ancestral state and sites with incorrect ancestral state. This approach has proved to be efficient for unbiased estimation of GC-biased gene conversion (Glémin *et al.* 2015), a weak selection-like process. Here, we also showed that we could capture the errors in the identification of ancestral state under a general distribution of fitness effects and, as apposed to the expectations of Galtier (2016), that the *r*_*i*_ parameters are not sufficient to correct for misidentification of ancestral state.

When using the divergence data in the inference, we corrected for mutations that were fixed in the sample but that were, in fact, polymorphic in the population. These mutations would incorrectly be counted into the divergence data. Our correction is different than the one used by Keightley and Eyre-Walker (2012), which is implemented in dfe-alpha. We found that this correction can lead dfe-alpha to predict values of *α* that are extreme, both on the positive and negative side. Our approach showed a much more consistent behavior throughout the simulations.

One drawback of the method presented here is that as the sample size n increases, so does the number of required r; parameters. Estimating too many parameters could lead to numerical difficulties in finding the optimum. One might expect that mutations present in; copies could be distorted to similar extent as mutations that are present in *i* - 1 or *i* + 1 copies. Using this, the number of *r*_*i*_ parameters could be reduced by allowing different consecutive polymorphism counts to share the same r parameter. A model selection procedure, via LRT or AIC, can then be used to decide on the most adequate grouping of the *r* parameters.

Similar to the *r*_*i*_ parameters, both our approach and methods that use probabilistic modeling to account for the identification of ancestral state rely on that the same process applies to both neutral and selected sites. This is probably not the case, as one could expect that the error in the identification of the ancestral state is different for the sites that are under selection. Theoretically, the neutral and selected sites could each have their own *ϵ*, but this would most likely not be identifiable. Nonetheless, it would be useful to investigate how robust the inference is when neutral and selected sites have different errors in the identification of the ancestral state. One could also put more effort in reducing the misidentification error when obtaining the unfolded from the folded SFS. Such an approach is pursued by Keightley *et al.* (2016), where the unfolded SFS is obtained by relying on two, instead of one, outgroup populations.

All methods that estimate the DFE require an a priori strict division of sites into neutral and selected classes. This is needed to disentangle the effects of selection from other forces, such as demography and misidentification of the ancestral state. It is expected that real data violates this assumptions, and it is not known to how extent this biases the inference. Similar to the *ϵ*, one could add a contamination error, *ϵ*_con_, with which, the observed neutral data would be modeled as a mixture of truly neutral sites and selected sites. However, this parameter would not be identifiable. It would though be interesting to investigate to what extent violations of this assumption bias the inference.

Throughout this paper, we used LRTs for model testing. However, inferences with or without divergence data are not comparable through LRT or AIC, or any other similar method (as the data are different). A goodness of fit test could be developed, that would investigate how closely the predicted SFS matches the observed one. This could then be used to decide if divergence data should be used in the inference or not.

Here, we assumed that selection is additive, where fitness of the heterozygote and derived homozygote are 1 + *s* and 1 + 2*s*, and the selection coefficient s is fixed in time. This assumption is made by most methods that infer the DFE and *α*, though some approaches exist for modeling arbitrary dominance or potentially temporal variation/fluctuations in selection coefficients. Williamson *et al.* (2004), Huerta-Sanchez *et al.* (2008) and Gossmann *et al.* (2014) pursue this in more details, illustrating the need for future development accounting for other types of selection regimes.

Our general approach can be applied to a wide range of species where the amount and impact of beneficial mutations on patterns of polymorphism and divergence varies widely (as uncovered by Galtier (2016)). Our method allows to accurately detect if beneficial mutations are present in the data, and a LRT can be used for model reduction, to let the data decide if a full or strictly deleterious DFE should be inferred. Importantly, we also show that estimating a full DFE, and thus learning about the property of beneficial mutations and expected amounts of adaptive substitution, is possible without relying on divergence data.

### Availability

The source code is available upon request from PT.

## Acknowledgments

We would like to thank Nicolas Galtier for useful early discussions and comments on the manuscript. This work has been supported by the European Research Council under the European Union’s Seventh Framework Program (FP7/20072013, ERC grant number 311341).

